# Individual differences in successful self-regulation of the dopaminergic midbrain

**DOI:** 10.1101/863639

**Authors:** Lydia Hellrung, Matthias Kirschner, James Sulzer, Ronald Sladky, Frank Scharnowski, Marcus Herdener, Philippe N. Tobler

**Author notes:** **<u>Corresponding author:</u>** Lydia Hellrung, Laboratory for Social and Neural Systems Research, Department of Economics, University of Zurich, Blümlisalpstrasse 10, 8006 Zurich, Switzerland.

## Abstract

The dopaminergic midbrain is associated with brain functions, such as reinforcement learning, motivation and decision-making that are often disturbed in neuropsychiatric disease. Previous research has shown that activity in the dopaminergic midbrain can be endogenously modulated via neurofeedback, suggesting potential for non-pharmacological interventions. However, the robustness of endogenous modulation, a requirement for clinical translation, is unclear. Here, we examined how self-modulation capability relates to regulation transfer. Moreover, to elucidate potential mechanisms underlying successful self-regulation, we studied individual prediction error coding, and, during an independent monetary incentive delay (MID) task, individual reward sensitivity. Fifty-nine participants underwent neurofeedback training either in a veridical or inverted feedback group. Successful self-regulation was associated with post-training activity within the cognitive control network and accompanied by decreasing prefrontal prediction error signals and increased prefrontal reward sensitivity in the MID task. The correlative link of dopaminergic self-regulation with individual differences in prefrontal prediction error and reward sensitivity suggests that reinforcement learning contributes to successful self-regulation. Our findings therefore provide new insights in the control of dopaminergic midbrain activity and pave the way to improve neurofeedback training in neuropsychiatric patients.

## 1 Introduction

The dopaminergic midbrain, including the ventral tegmental area (VTA) and substantia nigra (SN), plays a crucial role in reward processing, reinforcement learning^1–4^, motivation^5,6^, and decision-making^7^. Dysfunctions of the reward system have far-reaching consequences and are associated with the development of several severe psychiatric disease such as addiction^8^ and schizophrenia^9,10^. Despite decades of extensive neuroscience and imaging studies which have contributed to an impressive body of knowledge of normal and abnormal reward system function, the neural mechanisms controlling midbrain activity are still not fully understood^11^. One key issue that has received increasing attention is whether humans are able to cognitively control brain activity within the reward system. It has already been shown that both healthy controls^12,13^, and patients with cocaine addiction^14^ can learn to regulate SN/VTA activity during real-time functional magnetic resonance imaging (rt-fMRI) neurofeedback training. However, the outcome of primary interest in neurofeedback training is a transfer beyond training itself, i.e., the ability to regulate activity also after training and without feedback. Such transfer is critical for clinical applications, including those involving disorders of the reward system^15^. While MacInnes and colleagues^13^ observed significant neural transfer effects in the form of increased neural activity and connectivity of the VTA during transfer on the group level, the other two studies revealed high between-subject variance in this self-regulation success. The purpose of this work is to determine how variance arises, how individuals with successful transfer effects differ from individuals without transfer effects, and whether activity in brain regions other than the VTA characterize individuals with successful transfer. We addressed these issues by combining data from two previous rt-fMRI neurofeedback studies^12,14^ and pursuing three aims.

1. Our first goal was to characterize individual differences in the degree of successful transfer of SN/VTA self-regulation and thereby differentiate regulators from non-regulators. Individual differences in regulation success and high variability of transfer effects arises also in other neurofeedback modalities such as electroencephalography (EEG) and are often neglected^16^. For rt-fMRI neurofeedback control, neural activity in the cognitive (or executive) control network may play an important role especially when performing a demanding task such as imagery^17^. Therefore, and based on the known direct and indirect connections between prefrontal cortex and SN/VTA^18–21^ we hypothesize that successful transfer of SN/VTA regulation is associated with activation in brain regions that are part of the cognitive (executive) control network, especially prefrontal areas.
2. Our second goal was to determine whether the framework of (operant) associative learning can be used to explain neurofeedback training. In applications of the associative learning framework to neurofeedback^17,22^, the feedback provides a higher order reward and the chosen mental strategy is reinforced in proportion to the sign and magnitude of the feedback. At the beginning of the training, participants cannot predict which strategy will consistently lead to an up- or downregulation in brain activity within the target region. Therefore, if they use an adequate strategy, participants receive more reward than predicted corresponding to a positive prediction error. As a consequence, they would be more likely to repeat the strategy, expect higher feedback next time and gradually learn how to keep the feedback signal high. Accordingly, in regulators the size of the prediction error should gradually decrease as the expected feedback increasingly converges with the actual feedback. In contrast, for non-regulators and participants in a control group receiving unrelated or unstable feedback, the prediction errors would remain large and variable because these participants cannot learn any association between mental strategies and feedback. These straightforward implications of current theorizing about the mechanisms underlying neurofeedback remained largely untested (for a simulation study on the temporal dynamics of feedback: Oblak and colleagues^23^; for the correlation of BOLD with signal increase (‘success’) and decrease (‘failure’) during regulation: Radua and colleagues^24^. Here, we directly investigate the prediction error mechanism in regions that control the SN/VTA, which itself has been traditionally associated with the coding of reward prediction errors in both animal^2,25,26^ and human research^27,28^. Furthermore, the causal sufficiency of dopaminergic prediction error signals for learning has been reinforced by optogenetics^29,30^. Together, we hypothesize here that decreasing prediction error signals during neurofeedback learning are associated with successful self-regulation and transfer effects.
3. Our third aim was to relate individual differences in the ability to regulate the midbrain to characteristics of reward processing, in order to further distinguish regulators from non-regulators. Thus, we asked whether successful neurofeedback training, as measured by transfer effects, taps into general properties of the reward system. Given that adaptive reward processing characterizes the SN/VTA^1,31^ we used a variant of the monetary incentive delay (MID) task that captures differences in adaptive reward sensitivity between clinical and non-clinical populations^32^. Using this task, we tested the hypothesis that reward processing in regions that may control the dopaminergic midbrain is related to successful SN/VTA self-regulation.

In sum, to study individual differences in capability to gain control of the SN/VTA we used rt-fMRI neurofeedback training in healthy participants receiving either real feedback (veridical group) or inverted feedback (control group). We quantified the individual degree of successful transfer by comparing the individual post-training versus pre-training self-regulation capabilities. Moreover, we related individual differences in reward sensitivity to separately measured SN/VTA self-regulation success.

## 2 Methods

### 2.1 Participants

Fifty-nine right-handed participants (45 males, average age 28.25±5.25 years) underwent SN/VTA neurofeedback training. We analysed data from two independent projects, which used highly similar rt-fMRI paradigms, rt-fMRI software and scanner hardware. The first dataset^12^ comprised male participants, randomly assigned to one of two groups. The experimental group received veridical neurofeedback (N = 15), the control group received inverted neurofeedback (N = 16) as training signal. The second dataset^14^ comprised the healthy control participants (N=28, 14 males) of a project investigating also cocaine users (these data are not presented here). This group received veridical neurofeedback. A subset of the participants in the second dataset (N=25) also performed a variant of the monetary incentive delay (MID) task^32^. All participants provided written informed-consent and received compensation for their participation. The Zurich cantonal ethics committee approved these studies in accordance with the Human Subjects Guidelines of the Declaration of Helsinki.

### 2.2 Experimental setup and neuroimaging

All participants underwent neuroimaging in a Philips Achieva 3T magnetic resonance (MR) scanner using an eight channel SENSE head coil (Philips, Best, The Netherlands) either at the Laboratory for Social and Neural Systems Research Zurich (SNS Lab, Study 1) or the MR Center of the Psychiatric Hospital of the University of Zurich (Study 2). First, we acquired anatomical images (Study1: gradient echo T1-weighted sequence in 301 sagittal plane slices of 250 × 250 mm^2^ resulting in 1.1 mm^3^ voxels; Study2: spin-echo T2-weighted sequence with 70 sagittal plane slices of 230 × 184 mm^2^ resulting in 0.57 × 0.72 × 2 mm^3^ voxel size) prior to neurofeedback training and loaded them into BrainVoyager QX v2.3 (Brain Innovation, Maastricht, The Netherlands) to identify SN/VTA as target region (see 2.4 for details). To acquire functional data, we used 27 ascending transversal slices in a gradient echo T2*-weighted whole brain echo-planar image sequence in both studies. The in-plane resolution was 2 × 2 mm^2^, 3 mm slice thickness and 1.1 mm gap width over a field of view of 220 × 220 mm2, a TR/TE of 2000/35 ms and a flip angle of 82°. Slices were aligned with the anterior–posterior commissure and then tilted by 15°. Functional images were converted from Philips par/rec data format to ANALYZE and exported in real-time to the external analysis computer via the DRIN software library provided by Philips. This external computer ran Turbo BrainVoyager v3.0 (TBV – Brain Innovation, Maastricht, The Netherlands) to extract the BOLD signal from the images and calculate the neural activation for the feedback signal. The visual feedback signal was presented using custom-made software with Visual Studio 2008 (Microsoft, Redmond, WA, USA) through either a mirror mounted at the rear end of the scanner bore (Study 1) or through MR compatible goggles (Study 2).

### 2.3 Neurofeedback paradigm

The participants were instructed that their goal was to control a reward-related region-of-interest in their brains by imagining rewarding stimuli, actions, or events. We have previously shown that reward imagination activates SN/VTA with conventional fMRI^33^. Prior to scanning, we provided examples of such rewards, including palatable food items, motivating achievements, positive experiences with friends and family, favourite leisure activity or romantic imagery. We encouraged participants to use these different rewards as potential strategies for upregulating reward-related activity during the cue ‘Happy Time!’, here referred to as IMAGINE_REWARD condition. In contrast, during the cue ‘Rest’ (here referred to as REST condition), participants were asked to perform neutral imagery, such as mental calculation to reduce reward-related activity. In both conditions, real-time SN/VTA BOLD signal was continuously fed back to the participant visually with a smiley vertically translating proportional to the signal (Figure 1). Prior to training, participants were familiarized with the 5s delay of the hemodynamic response affecting the display of the feedback and were asked not to move or change their breathing during the neurofeedback training.

**Figure 1.**
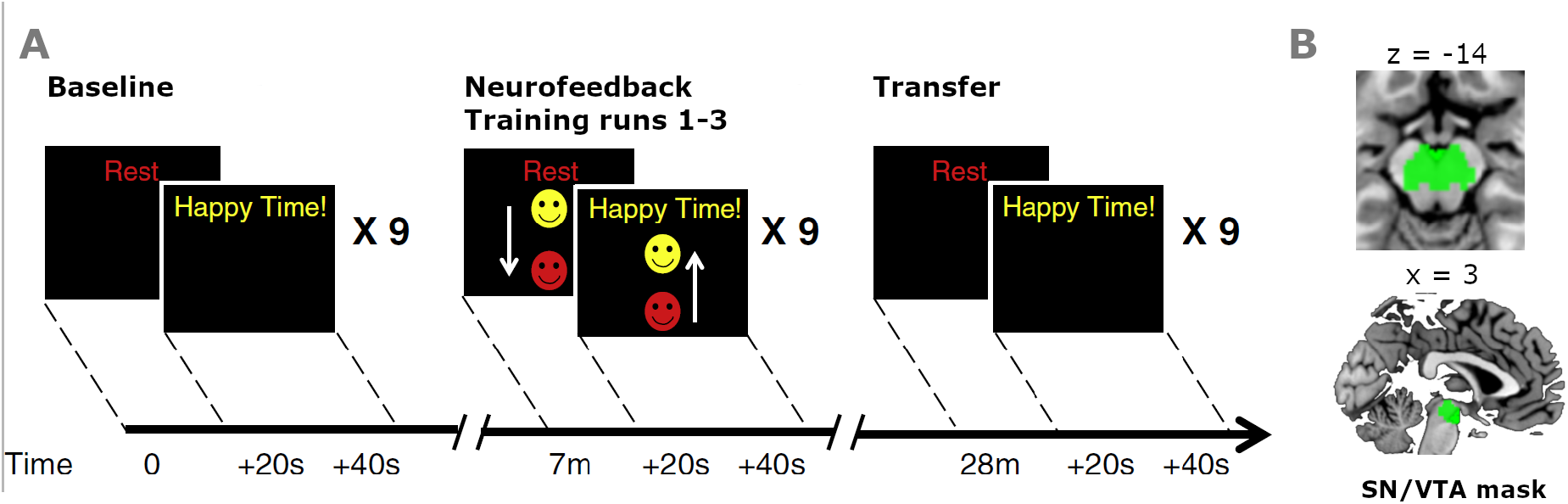
Neurofeedback paradigm: (A) All runs consisted of alternating blocks of REST and IMAGINE_REWARD conditions, with each block lasting 20 s. The regulation conditions (REST, IMAGINE_REWARD) were indicated by words (‘Rest’ or ‘Happy Time!’) and the feedback presented as moving smiley face during neurofeedback training runs. The baseline and transfer runs comprised no feedback. The SN/VTA signal difference from these runs served to quantify the degree of regulation transfer (DRT) as (SN/VTA_BOLD_{IMAGINE_REWARD, Transfer}_ − SN/VTA_BOLD_{REST, Transfer}_) − (SN/VTA_BOLD_{IMAGINE_REWARD, Baseline}_ − SN/VTA_BOLD_{REST, Baseline}_). (B) Post-processed SN/VTA signal was extracted from the probabilistic atlas mask^34^.

Each neurofeedback session comprised: a pre-training imagery baseline run without any feedback, three (Study 1) or two (Study 2) training runs during which neurofeedback was presented (as Study 2 also investigated patients, training was limited to two runs), and a transfer run (i.e., without feedback). Each of these runs comprised nine blocks of IMAGINE_REWARD and REST conditions, each lasting 20 s. To determine the current level of the feedback signal we used the average of the last five volumes of the previous REST condition as reference value and employed a moving average of the previous three volumes to reduce noise. In the veridical feedback group, the smiley moved up with increasing percent signal change of SN/VTA BOLD signal and changed colour from red to yellow (Fig. 1 A). In the inverted feedback group, the smiley moved up and turned yellow with a decreasing SN/VTA BOLD signal.

### 2.4 Region-of-interest SN/VTA

In both studies, the target region for neurofeedback, i.e. the substantia nigra (SN) and ventral tegmental area (VTA), was structurally identified using individual anatomical scans. Since the individual mask definition slightly differed between Study 1 and 2 (T1-weighted scans in Study 1 and T2-weighted scans in Study 2), we used an independent mask for our post-hoc analysis. By this, we can control for individual differences between experimenter ROI selection strategies, to avoid interpolation confounds due to warping by normalization and use a reliable seed region for functional connectivity analysis. Specifically, we used the probabilistic mask of the SN and VTA as defined by^35^, which is based on a large sample (148 datasets) and available on https://www.adcocklab.org/neuroimaging-tools (download August 2018). Figure 1B illustrates this mask within the brain. From this mask image, we extracted and averaged SN/VTA activity for each participant using custom-made scripts in Matlab R2016b.

### 2.5 Degree of regulation transfer (DRT)

We assessed the effects of individual differences in performance to characterise participants on a continuous regulation scale. The measure of successful self-regulation was defined as individual degree of regulation transfer (DRT), i.e. as the condition-specific SN/VTA signal difference between post-training (Transfer) and pre-training (Baseline) runs:

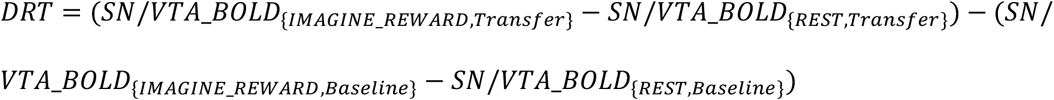

Thus, a positive DRT corresponds to a relative increase in post-training SN/VTA BOLD activity compared to pre-training SN/VTA BOLD activity for the contrast IMAGINE_REWARD minus REST. It is essential to note that during these two runs (pre-training baseline, post-training transfer) no neurofeedback was presented. This definition ensures comparability between participants in the different intervention groups, while the perception and processing of the feedback signal during the training runs might be different and influencing the SN/VTA signal itself.

To achieve positive transfer effects, participants had to apply what they had learned during training runs. We therefore asked whether DRT is related to SN/VTA activity during the training runs by calculating the correlation between them. For this, we used the slope of SN/VTA signal change increase over training time in Spearman’s correlations for the intervention and control group.

#### DRT distributions

To investigate potential group differences in DRT, we transferred the extracted data to R (R-project R3.4.1). Using an ANOVA and a non-parametric Kruskal-Wallis test, we tested for differences of the mean between the three groups (i.e. the two groups receiving veridical feedback in Studies 1 and 2 and the control group receiving inverted feedback in Study 1).

#### DRT in fMRI analysis

The DRT measure served to investigate the individual differences in successful transfer at the whole brain level. In particular, we were interested to identify regions that were positively associated with DRT and thus potentially contribute to regulation of the SN/VTA. For this analysis, we entered mean-centered individual DRT levels in all fMRI second level statistical models (see 2.8). We excluded SN/VTA from all analyses to avoid any circularity.

#### Spatial specificity control analysis

To investigate the spatial specificity of our analysis of dopaminergic midbrain regulation, we performed the same whole brain analysis as described above for SN/VTA with a different ROI. Specifically, we used the neighboring brain region of the parahippocampus (Supplemental Material). In keeping with specificity, this control analysis revealed little commonality (limited to the cerebellum and temporal gyrus) with the SN/VTA analysis (Figure S4 and Table S8).

### 2.6 MID Task

In addition to the neurofeedback training, the participants in Study 2 (N=25) performed a MID task that captures differences in adaptive reward sensitivity. In every trial of the MID task^32,36,37^ first one of three cues appeared (Fig. S1). One cue was associated with large reward (ranging from 0 to 2.00 CHF), one cue with small reward (0 to 0.40 CHF) and one cue with no reward. After a delay of 2.5 to 3 s, participants had to identify an outlier from three circles by pressing one of three buttons as quickly as possible. Depending on the cue, their response time and the correctness of the answer, participants gained an amount of money. Importantly, the use of large and small reward ranges enables investigation of individual differences not only in general reward sensitivity but also in how well the reward system adapts to different reward distributions, so-called adaptive reward coding^32^.

### 2.7 MR Data pre-processing

We despiked the functional data using AFNI toolbox (National Institute of Mental Health; http://afni.nimh.nih.gov/afni). To account for differences in echo-planar-image (EPI) slice acquisition times we employed temporal interpolation of the MR signal, shifting the signal of the misaligned slices to the first slice^38^ using FSL 5 (FMRIB Software Library, Analysis Group, FMRIB, Oxford, http://fsl.fmrib.ox.ac.uk). Furthermore, data were bias-field corrected using ANTs (Advanced Normalization Tools; http://stnava.github.io/ANTs), realigned using FSL 5, normalized to standard Montreal Imaging Institute (MNI) space using ANTs in combination with a custom scanner-specific EPI-template resulting in a 1.5 mm^3^ isotropic resolution and finally smoothed with a 6 mm full-width-half-maximum Gaussian kernel using FSL 5.

The spatial specificity control analyses (Figure S4 and Table S8) suggest that the findings reported here are not due to common physiological noise. To more directly account for noise, we additionally acquired physiological data in a subsample of participants. In the available subsample, neither changes in heart rate variability nor respiration were significantly correlated with VTA/SN activation during reward imagination (see details in Kirschner et al.^39^, Supplemental Material Table S1, Figure S1). Here, we also used an image-based correction to account for physiological artefacts in all participants. Since physiological artefacts are most prominently present in CSF and white matter due to the absence of BOLD effects, pulsations of the ventricles, and proximity to the large brain arteries (e.g., circle of Willis), we decided to use an established preprocessing procedure based on a principal component analysis (PCA) approach^40,41^. Specifically, we calculated the global mean and the first 6 components of a temporal principal component analysis on the cerebrospinal fluid and white matter signal. These 6 components were used as noise regressors in the first level statistics (see 2.8) in addition to the 6 motion parameters. Along with the pre-processing of the fMRI data, the SN/VTA mask used as ROI for the analysis was resliced into the dimensions of the functional data using SPM 12 (v6906, Wellcome Trust Centre for Neuroimaging, UCL, London, UK; http://www.fil.ion.ucl.ac.uk/spm/software/spm12/) within Matlab R2016b (Mathworks, Sherborn, MA, USA).

### 2.8 MR Data analysis

For all of the following analyses, we used the toolbox SPM 12 (v6906) within Matlab R2016b. All figures were created using bspmview v.20161108^42^ and ggplot2 within R 3.4.1. All group-level analysis included an additional covariate for the dataset to account for potential global signal differences between studies.

#### Post-training effects: Correlation with DRT in veridical and inverted feedback group (aim 1)

The first question of this study asked whether the individual degree of successful neurofeedback transfer is associated with individual differences in the cognitive control network. To answer this question, we conducted a general linear model (GLM) on the single subject level including one block-wise regressor for the IMAGINE_REWARD condition and one for the REST condition with 190 timesteps (each condition comprised 9 onsets and lasted 20 s) for each of the four runs separately. Additionally, we modelled the first 5 TRs of every run as nuisance regressor and added also motion and physiological artefact regressors (see section 2.7) in the design matrix. In total the GLM consisted of fifteen regressors. We formed the contrast IMAGINE_REWARD-REST and compared it between Transfer and Baseline runs, i.e. (IMAGINE_REWARD-REST)_Transfer_ − (IMAGINE_REWARD-REST)_Baseline_. At the group level, we tested for correlation of the SN/VTA-derived DRT with this contrast in a one-sample t-test. We ran these analyses in all voxels other than the SN/VTA and separately for both the veridical and inverted feedback groups. To test for common and separate activity between the groups, we performed conjunction and disjunction analyses over the two group maps. Additionally, we performed a two-sample t-test group comparison analysis to identify significant group differences. To identify activity within the cognitive control network, we used a cognitive control template based on the coordinates from a meta-analysis^43^. We created this template with fslmaths and spheres of 15 mm around all coordinates from the meta-analysis. In table S1 we identify regions of the cognitive control network where transfer success correlates with DRT within the template. For statistical maps, we used FWE-corrected cluster level threshold with p < 0.05 (cluster extent of 230 voxel) based on whole brain statistics p < 0.001. In addition, to test the functional specificity of our results, we performed a meta-analytic functional decoding analysis using the Neurosynth database (www.neurosynth.org). This relates the neural signatures of the cognitive control decoding network to other task-related neural patterns (Fig. S2).

#### Prediction error coding analysis during NF training (aim 2)

The second question of the study asked whether successful neurofeedback performance was associated with a reduction in prediction error during the training runs as captured by a classic reinforcement learning framework. To address this issue, we determined the temporal difference of the feedback signal (i.e., the change in height of the smiley) as proxy for the prediction error signal. Specifically, for the neurofeedback training runs we constructed a GLM that replaced the block-level (IMAGINE_REWARD and REST) regressors with corresponding event-level regressors that modelled every TR and that we parametrically modulated with a time-resolved continuous prediction error (PE) term. This PE term was defined as difference between the current and the previous TR within the SN/VTA mask, i.e. (BOLD_SN/VTA_t_- BOLD_SN/VTA_t−1_; accordingly, in the upregulation condition the parametric modulator corresponded to IMAGINE_REWARD_t_ ‒IMAGINE_REWARD_t−1_). To investigate if the prediction error decreases over time, we used the difference (parametric modulator PE (run 2) – parametric modulator PE (run 1), i.e. PE coding in neurofeedback training run 2 minus neurofeedback training run 1 (Figure 1A). This difference should become negative as prediction errors decrease with learning. On the group level, we correlated this contrast (difference in PE coding run2 – PE coding run1) with the DRT measure in a one-sample t-test to test for associations between a decrease in prediction error coding and successful self-regulation.

The results of this analysis, showing prediction error coding in the dorsolateral prefrontal cortex (dlPFC), inspired a functional connectivity analysis. Specifically, we investigated the functional impact of the dlPFC prediction error signal on the SN/VTA using a psychophysiological interaction analysis using the gPPI v13 Toolbox^44^ based on the MNI coordinate of dlPFC (x=40, y=10, z=38) with a 5 mm sphere as seed region. We added activity from this seed region as physiological regressor to the original GLM and interacted it with both the IMAGINE_REWARD and REST regressors to form interaction regressors. Functional connectivity was calculated by contrasting the interaction terms IMAGINE_REWARD-REST between second and first neurofeedback training run. We then correlated this contrast with DRT. The results were focused to the SN/VTA region as target. For statistical maps, we used a whole-brain threshold of p < 0.001 (20 voxel extent).

#### Relation between DRT and reward sensitivity in the MID Task (aim 3)

To address the third aim of the study, we investigated the relationship between reward processing in the MID task and the capacity to successfully regulate the SN/VTA in the neurofeedback experiment. In particular, we considered two contrasts in the MID task (1) general reward sensitivity, defined as the sum of parametric modulators: small plus large reward (2) adaptive reward coding, defined as the difference between parametric modulators: small minus large reward. Again, we used correlation analysis at the group level to determine whether these two contrasts are related with individual SN/VTA transfer success (DRT) in the neurofeedback task. Moreover, to assess the commonalities of the neural activities in these different tasks, we performed a conjunction analysis of contrasts (1), (2) and the correlation of transfer-activity with DRT (see 2.8). For statistical maps, we used a whole-brain threshold of p < 0.001 (20 voxel extent due to conjunction).

### 2.9 Additional behavioral measurements

#### Strategies

All participants were introduced to five example strategies (see 2.3) that they could use to upregulate brain activity but also free to use their own strategies. At the end of the experiment, participants filled in a custom-made questionnaire on the strategies they used. To compare strategies between the groups, we used a χ2-test to assess differences in the distribution of strategy usage. We did not observe any significant group differences in strategy use (p = .9), and therefore did not consider this measurement in any further analysis.

#### Personality measures

To investigate whether individual differences in behavior and personality were associated with individual differences in DRT, Study 2 measured: (1) Smoking status in number of cigarettes per day; (2) verbal IQ as determined by the Multiple Word Test (MWT^45^); (3) Positive and Negative Affect Score (PANAS) in the German version^46^; (4) attentional and nonplanning subscores of the Barratt Impulsivity Scale in the German version^47^. We tested for correlations with the DRT parameter using Pearson correlations. As none of these variables correlated significantly with the DRT parameter (all p > 0.5), we did not consider them further.

## 3 Results

### 3.1 No difference in degree of regulation transfer (DRT) across groups

We first evaluated the DRT measure and compared it between the three datasets. There were no significant differences across all three groups (mean DRT veridical group Study 1 = 0.01, mean DRT veridical group Study 2 = −0.02, mean DRT inverted group Study 1= −0.05; anova testing F(2, 56) = 0.13; non-parametric Kruskal-Wallis testing H(2) = 0.39, p = 0.82; Fig. 1; see Supplemental Figure S5 for alternative illustration). Moreover, also the direct comparison between the two veridical groups was not significant (T(39) = −0.26, *p* = 0.8). Accordingly, we combined the two veridical groups for subsequent analyses. Importantly, our participants showed considerable variation in DRT, which allowed us to investigate the individual differences in brain activity accompanying more or less successful regulation of the SN/VTA through neurofeedback. Thus, the groups showed similar mean levels and considerable individual differences in self-regulation success.

### 3.2 Correlation of slopes between transfer and training only for intervention group

Next, we tested for differences between groups in the relationship of SN/VTA transfer as measured by DRT with signal change in SN/VTA during training. We found positive correlations between the slope of SN/VTA signal change during training period and DRT only for the veridical feedback group, but not for the control group (veridical group ρ = 0.62, p < 0.001; inverted group Rho = −0.3, p = 0.25). Although the comparability between training and transfer is limited due to the feedback signal processing, this indicates that only the veridical feedback group benefitted from the feedback. More importantly, within the veridical feedback group, particularly those individuals who were more successful at transfer also showed stronger upregulation during training.

### 3.3 Individual variation in transfer: DRT associated with cognitive control network in veridical and amygdala activity in inverted feedback group

#### 3.3.1 Veridical feedback group

We investigated whether individual levels of successful SN/VTA self-regulation (DRT) were associated with increased post-training activity compared to pre-training activity in other regions of the brain (IMAGINE_REWARD-REST)_Transfer_ − (IMAGINE_REWARD-REST)_Baseline_). This analysis revealed several areas consistently reported by neurofeedback studies (see Fig. 2 in the meta-analysis of Sitaram et al.^17^, including dorsolateral prefrontal cortex (dlPFC), anterior cingulate cortex (ACC), lateral occipital cortex (LOC), and thalamus (Figure 3A and Table 1). To formally test for a more general association with the cognitive control network, we applied a cognitive control network template from a meta-analysis^43^, which in addition revealed neural activity in precuneus and striatum (Fig. 3B for exemplary illustrations of dlPFC, ACC, temporal gyrus, and thalamus activity; Table S1 for full overview). Thus, regions of the cognitive control network showed transfer to the extent that neurofeedback training of the dopaminergic midbrain was successful.

**Table 1:** Correlation of transfer activity (IMAGINE_REWARD_transfer_ − REST_transfer_) − (IMAGINE_REWARD_baseline_ − REST_baseline_) with DRT in veridical feedback group (see Figure 3a). Table shows all local maxima separated by more than 20 mm; for all clusters, p < 0.05 FWE-corrected on cluster level; df = 40. Regions were labelled using the Harvard-Oxford atlas and/or the Anatomy Toolbox in parentheses; the activity in SN/VTA has been excluded from the table to avoid circularity; x,y,z = Montreal Neurological Institute (MNI) coordinates in the left-right, anterior-posterior, and inferior-superior dimensions, respectively.

| Region Label | # voxels | t-value | MNI Coordinates |  |  |
| --- | --- | --- | --- | --- | --- |
|  |  |  | x | y | z |
| Cingulate Gyrus, posterior division | 895 | 6.104 | -3 | -49 | 7 |
| Middle Frontal Gyrus<br>(dorsolateral prefrontal cortex) | 295 | 4.609 | 45 | 31 | 19 |
| Left Thalamus | 1281 | 4.472 | -9 | -24 | 10 |
| Temporal Occipital Fusiform Cortex | 715 | 5.858 | 32 | -46 | -22 |
| Lingual Gyrus (Right<br>Parahippocampal Gyrus) | 715 | 3.725 | 20 | -46 | -6 |
| Right Cerebral White Matter<br>(Right Hippocampus) | 298 | 5.300 | 20 | -18 | -10 |
| Left Hippocampus | 408 | 3.703 | -26 | -33 | -12 |
| Left Cerebral White Matter<br>(Left Middle Occipital Gyrus) | 693 | 5.056 | -36 | -73 | 31 |
| Left Cerebral White Matter<br>(Left Superior Medial Gyrus, Anterior<br>Cingulate Cortex) | 579 | 4.960 | -9 | 28 | 38 |
| Intracalcarine Cortex<br>(Right Lingual Gyrus) | 856 | 4.048 | 6 | -82 | 1 |
| Occipital Fusiform Gyrus | 474 | 4.928 | 29 | -66 | -12 |
| Middle Temporal Gyrus (Left Inferior<br>Temporal Gyrus) | 1028 | 4.913 | -62 | -37 | -16 |
| Middle Temporal Gyrus (Left Inferior<br>Temporal Gyrus) | 1028 | 4.315 | -59 | -57 | -3 |
| Occipital Fusiform Gyrus | 658 | 4.866 | -42 | -70 | -19 |
| Temporal Fusiform Cortex, posterior<br>division | 658 | 4.638 | -39 | -43 | -28 |
| Temporal Occipital Fusiform Cortex | 658 | 4.401 | -23 | -66 | -19 |
| Superior Frontal Gyrus | 310 | 4.736 | 6 | 2 | 80 |
| Central Opercular Cortex<br>(Left Superior Temporal Gyrus) | 309 | 4.636 | -59 | -16 | 16 |
| Left Cerebral White Matter<br>(Left inferior Frontal Gyrus) | 401 | 4.589 | -39 | 35 | -9 |
| Location not in atlas | 408 | 4.578 | -14 | -49 | -22 |
| Location not in atlas (Right<br>Paracentral Lobule) | 341 | 5.003 | 2 | -39 | 82 |
| Location not in atlas<br>(Left Cerebellum IV) | 856 | 4.947 | -6 | -66 | -15 |

**Figure 2.**
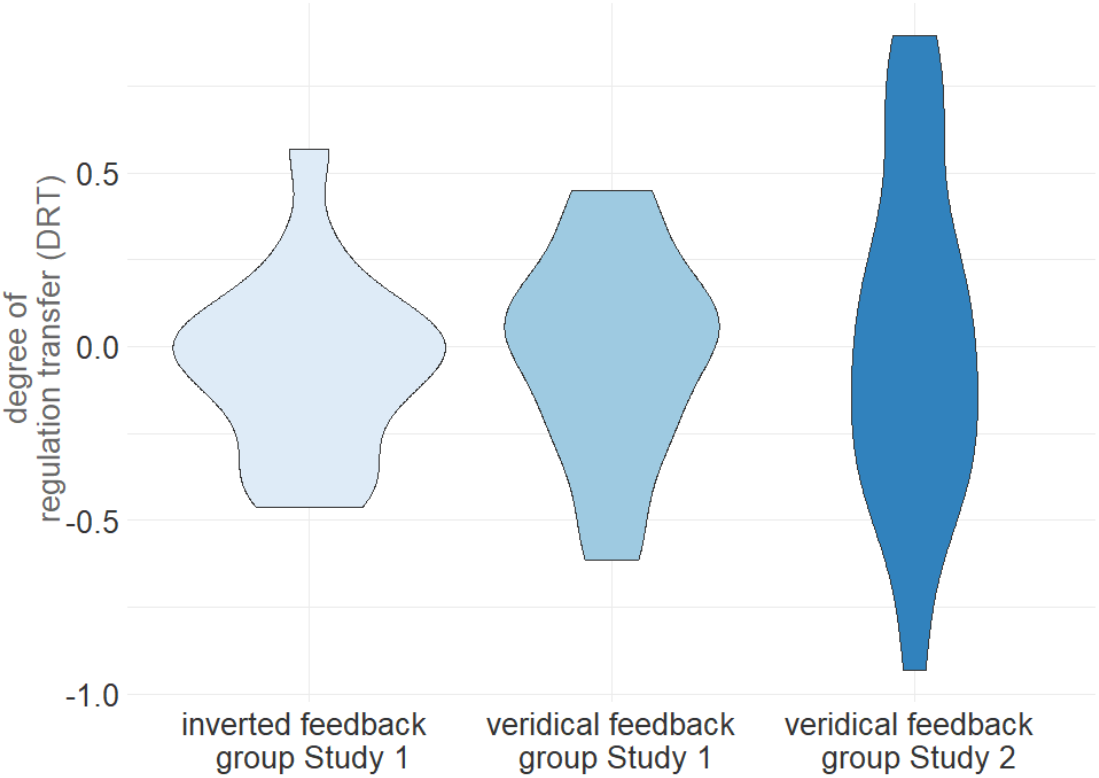
Distribution of DRT across groups. The DRT measure was distributed similarly in both groups receiving veridical feedback in Studies 1 and 2 and the control group receiving inverted feedback in Study Accordingly, we found no evidence supporting a main effect of feedback on transfer. However, DRT varied substantially across individuals, which motivated the analyses using the individual self-regulation success.

**Figure 3:**
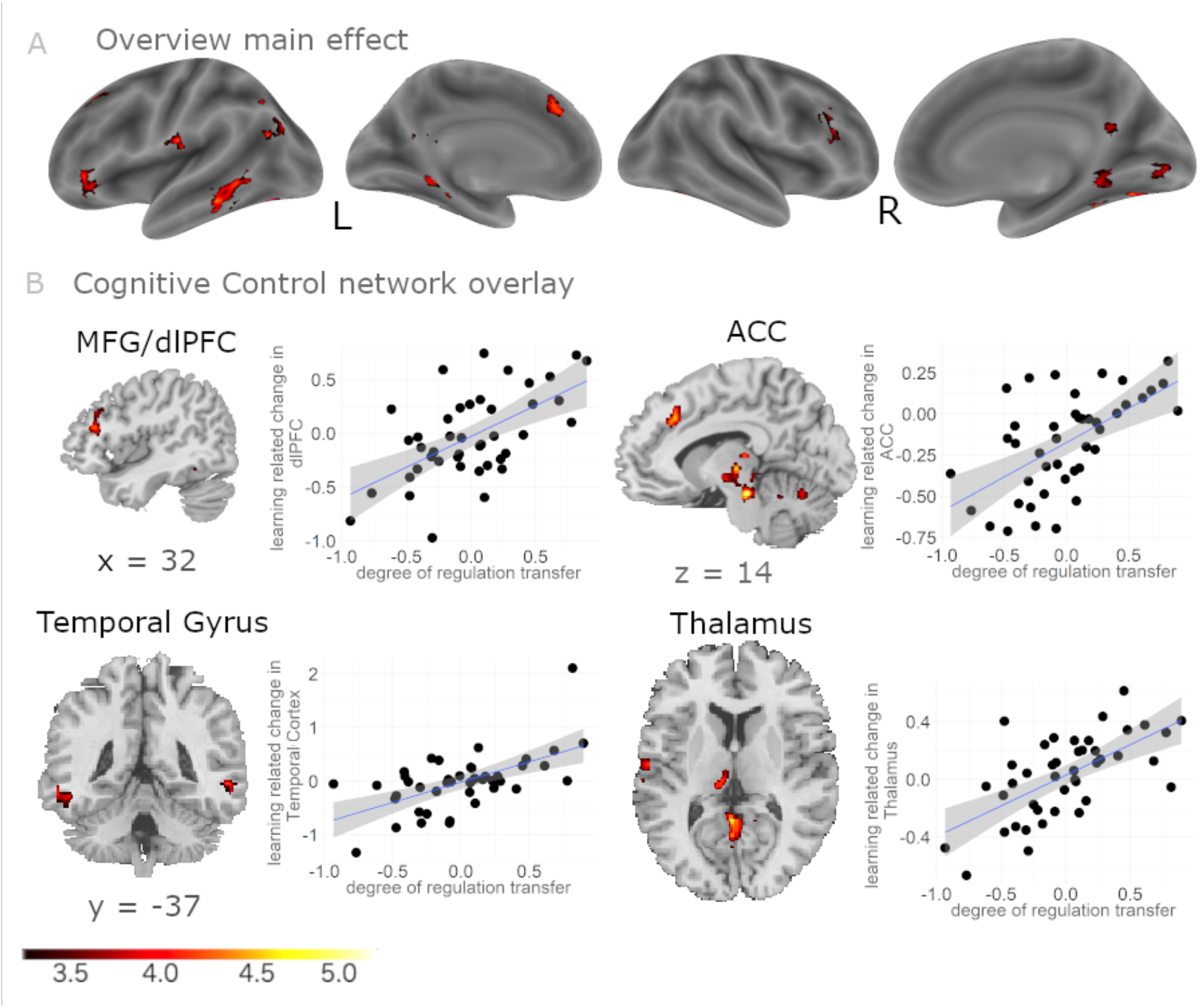
Correlation of DRT with transfer success after training in veridical feedback group: To investigate whole-brain neural activity correlating with successful SN/VTA self-regulation, we used DRT as measure of successful regulation of the SN/VTA and correlated it with the contrast (IMAGINE_REWARD_transfer_ − REST_transfer_) − (IMAGINE_REWARD_baseline_ − REST_baseline_) as measure of learning related change in neural activity in the rest of the brain. A) The analysis revealed task-specific correlations primarily within the cognitive control network (whole brain overview FWE-corrected with p < 0.05 on cluster level, projected to lateral and medial sagittal sections). B) Exemplary correlations within the cognitive control network have been depicted, here in MFG/dlPFC, ACC, Thalamus, and bilateral Temporal Gyrus, to illustrate the association between neural activity with DRT. The correlations are for illustration purposes only without further significance testing to avoid double dipping. The grey shaded area identifies 95 % confidence interval.

#### 3.3.2 Inverted feedback group

For the inverted feedback group, the same analysis resulted in partly distinct activations. In contrast to the veridical feedback group, left amygdala activity correlated significantly with DRT (Fig. 4 and Table S2). Importantly, activity in cognitive control areas reported above, such as dlPFC and ACC, was significantly weaker in inverted than veridical feedback groups (Table S3 for disjunction and direct statistical comparison). Together with the lack of correlation of DRT with SN/VTA signal change during training for the inverted feedback group, these findings suggest that cognitive control regions play a preferential role for successful transfer of SN/VTA self-regulation.

**Figure 4.**
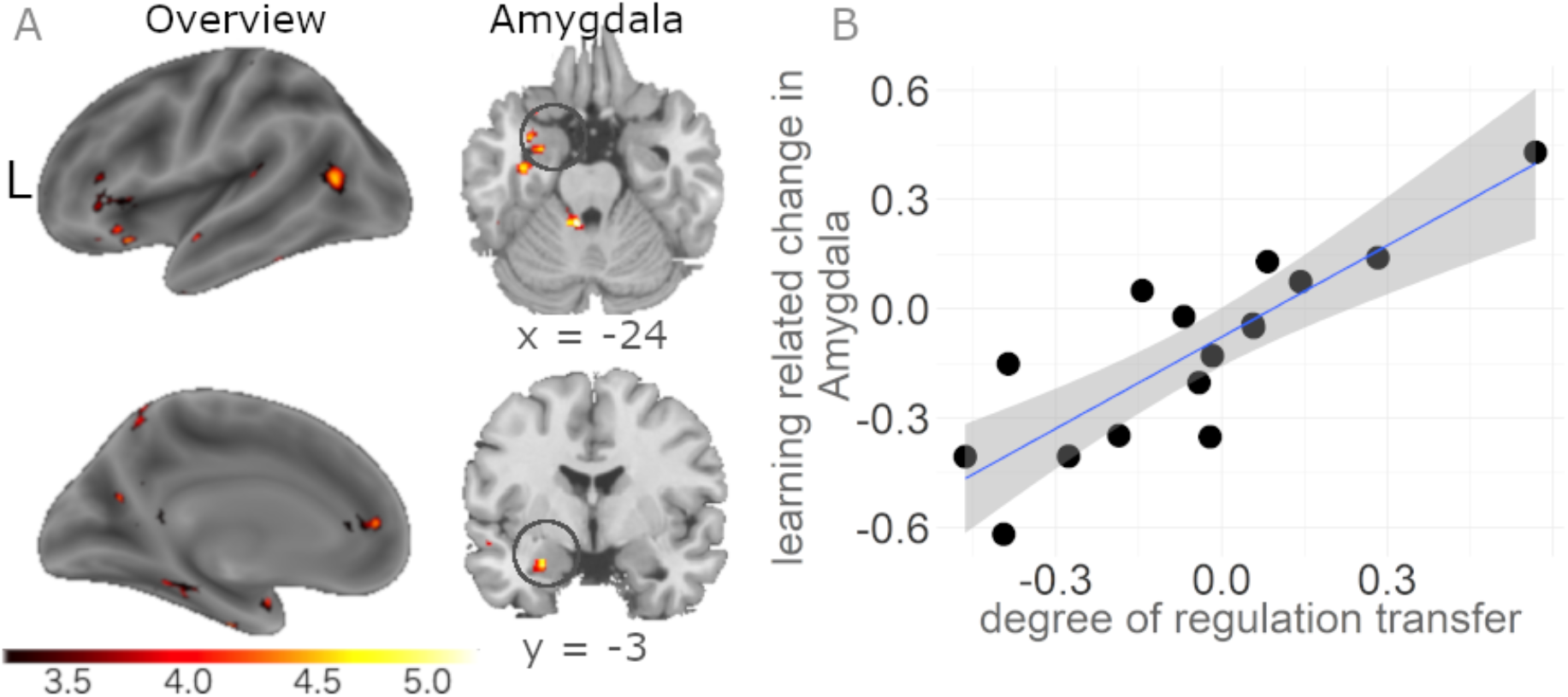
Correlation of DRT with transfer success after training in inverted feedback group: (A) Receiving inverted feedback resulted in a correlation between DRT as measure of regulation success and the contrast (IMAGINE_REWARD_transfer_ − REST_transfer_) − (IMAGINE_REWARD_baseline_ − REST_baseline_) as measure of learning related change in the amygdala (p < 0.001). This region was not observed in the veridical group. (B) The correlation depicts the positive association of neural activity in the amygdala with DRT. The plot is for illustration purposes only without further significance testing to avoid double dipping. The grey shaded area identifies the 95 % confidence interval.

We also tested for common activity in the two feedback groups using conjunction analysis. Similar to the veridical group, the inverted feedback group showed correlations between DRT and activity in the precuneus, middle temporal gyrus, insula, thalamus, and parahippocampal gyrus (Table S4). These common areas appear to reflect non-specific regulation activity and may be associated with memory and introspection processes.

### 3.4 Reinforcement learning: DLPFC prediction error coding during neurofeedback training correlates with DRT

To investigate whether reinforcement learning mechanisms contribute to successful neurofeedback transfer, we tested for the temporal differences in the feedback signal as proxy for the prediction error signal during the training runs. We reasoned that prediction error activity should decrease from early to late phases of neurofeedback training for successful regulators. At any time during neurofeedback training, participants needed to come up with their own predictions of the upcoming feedback signal and compare the predictions with actual feedback at the next time point. Similarly, in temporal difference learning models, prediction errors are calculated at each moment in time^48^. Therefore, we operationalized prediction error by subtracting the immediately preceding SN/VTA activity (prediction) from the present SN/VTA activity (outcome). Specifically, we tested for a negative correlation of DRT with the difference in prediction error coding signals between late and early training. In other words, only for participants with high DRT we expected to observe a decrease of prediction error signal over the course of the neurofeedback training. We found such gradually decreasing prediction error signals in dlPFC (Fig. 5 and Table S5). To interrogate the finding in detail, we also analysed the two neurofeedback training runs separately. This analysis confirmed that only successful regulators showed less pronounced dlPFC coding of prediction error in late compared to early training (see Fig. S3 for run-wise PE coding in dlPFC). Importantly, it should be noted that this decrease of error signals in dlPFC is related to the individual DRT levels. The basic contrast of prediction error coding, i.e. without correlation to DRT, revealed striatal activity.

**Figure 5:**
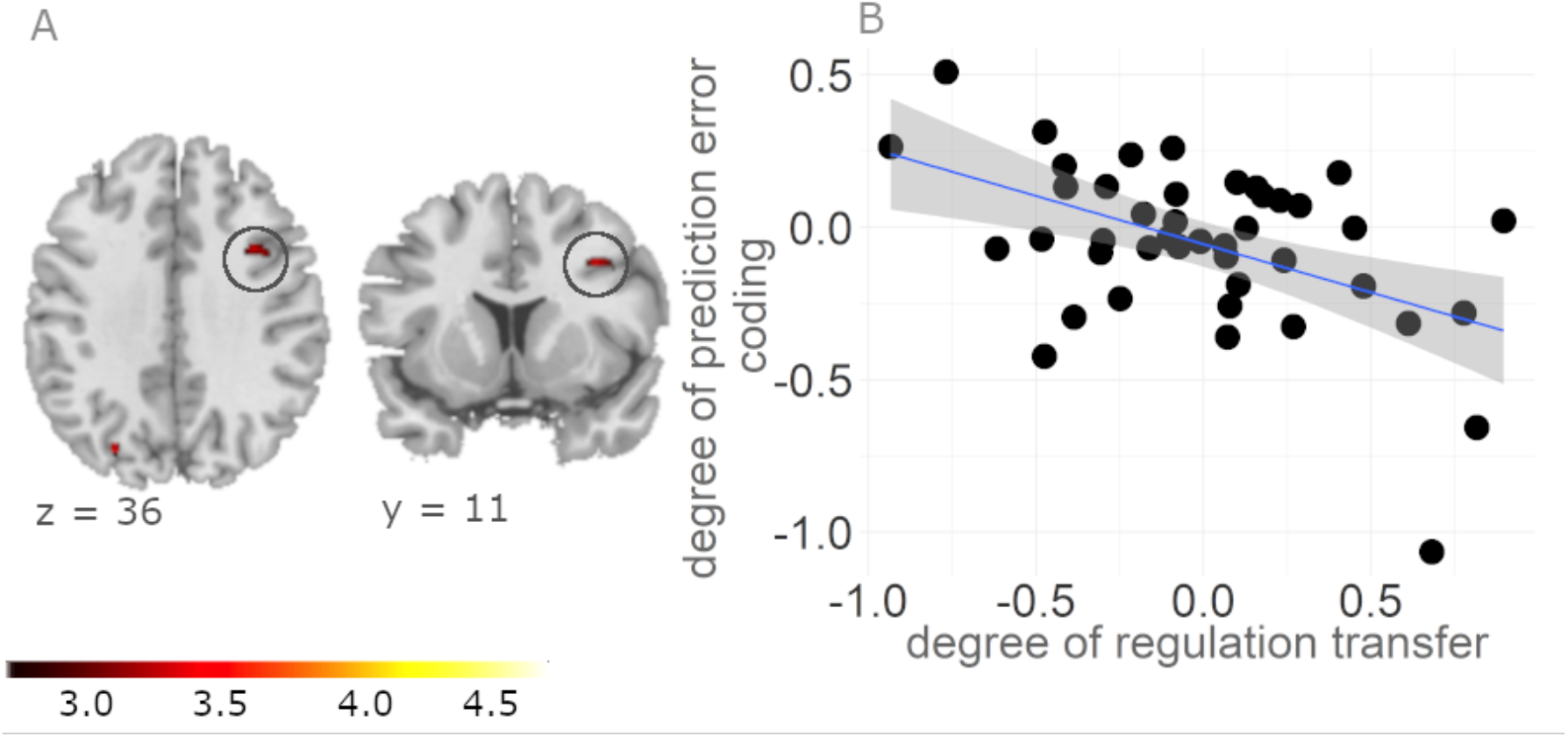
Prediction error coding in dlPFC decreases during NF training in participants with successful SN/VTA self-regulation: (A) The neural prediction error signal, corresponding to the temporal difference between the current and immediately preceding feedback activity from the SN/VTA decreased with ongoing feedback training (i.e, the difference between the last and first run) within dlPFC more strongly in individuals with higher DRT (p < 0.001). This finding is consistent with reinforcement learning theories, according to which prediction errors decrease as learning progresses. By extension, a reinforcement learning framework can explain successful neurofeedback training. (B) The plot depicts the differences in prediction error signals in dlPFC between the last and first training for every participant. This shows that the individual degree of regulation success statistically relates to the decrease in prediction error coding over training. The plot is for illustration purposes only without further significance testing to avoid double dipping. The grey shaded area identifies the 95 % confidence interval.

### 3.5 Learning-related functional coupling of DLPFC with SN/VTA

Following on from our finding of decreasing prediction error coding in dlPFC being related to individual success of regulating the dopaminergic midbrain, we performed a functional connectivity analysis to investigate whether the identified dlPFC region communicates with the SN/VTA region our participants aimed to regulate. Thus, we used the dlPFC region showing decreasing prediction error coding during training particularly in successful regulators as a seed region and investigated the coupling to the SN/VTA. Functional connectivity between the two regions increased with transfer success (Fig. 6; (t(40) = 3.79, cluster extent = 16, MNI × = −2, y = −16, z = −15). In other words, DRT and dlPFC to SN/VTA connectivity correlated positively. Note that this correlation of DRT with dlPFC-SN/VTA connectivity was task-related as it was enhanced during IMAGINE_REWARD relative to REST (which served as psychological regressor) and independent of SN/VTA activity.

**Figure 6.**
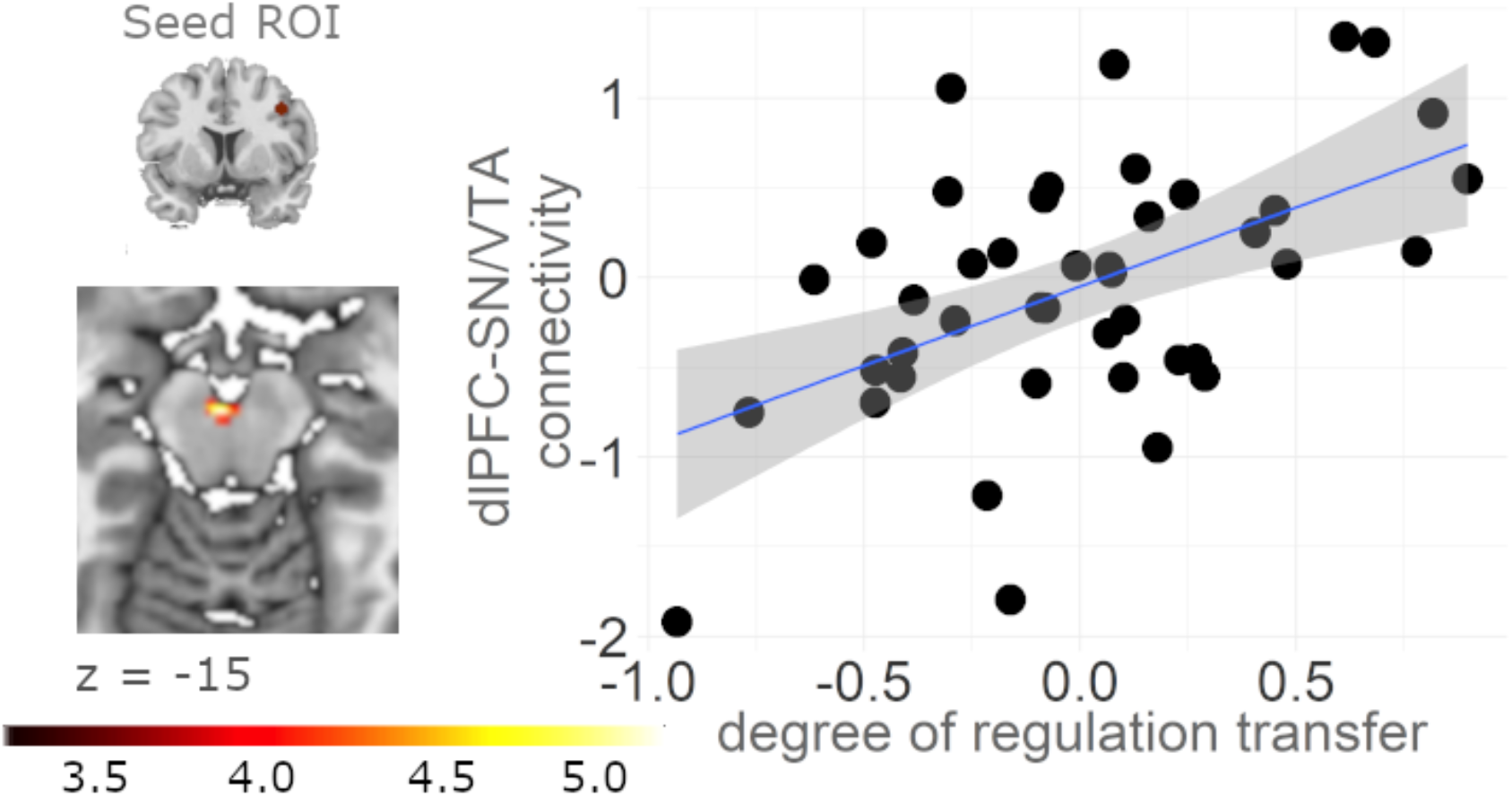
Functional connectivity between dlPFC and SN/VTA correlates with transfer success: (A) A functional connectivity analysis based on the prediction error coding seed region in the dlPFC (MNI coordinate 40, 10, 38, 5 cm sphere) revealed that connectivity with the SN/VTA correlated positively with success of neurofeedback training (p < 0.001). (B) DRT increased with increasing connectivity between dlPFC and SN/VTA during IMAGINE_REWARD vs. REST in neurofeedback training runs. Thus, dlPFC appears to communicate with SN/VTA in proportion to the degree to which neurofeedback training is successful. (C) The correlation plot depicts connectivity between dlPFC and SN/VTA with DRT. The plot is for illustration purposes only without further significance testing to avoid double dipping. The grey shaded area identifies the 95 % confidence interval.

### 3.6 Individual differences in dlPFC reward sensitivity during MID task correlate with regulation success

In Study 2 we used the MID task to independently measure reward sensitivity and the capability to adapt to different reward contexts^32^. We asked whether individual measures of reward processing (measured with parametric and adaptive coding of reward related BOLD activity) are related to individual success in regulating the SN/VTA. Specifically, we tested for correlations between DRT and (i) MID reward sensitivity (sum of small and large reward parametric modulators) and (ii) MID adaptive reward coding (difference of small minus large reward parametric modulators). These two correlations both identified dlPFC (Fig. 7A). Moreover, a conjunction of these two correlations with the correlation between DRT and the contrast (IMAGINE_REWARD_transfer_ − REST_transfer_) − (IMAGINE_REWARD_baseline_ − REST_baseline_) outside SN/VTA revealed common neural activity in the dlPFC (center at MNI x = 40, y = 10, z = 38; Fig. 7 and Table S6). Thus, the more successful individuals were at self-regulating SN/VTA as a result of neurofeedback training, the more sensitive they were to reward and the more strongly they adapted to different reward contexts in the MID task.

**Figure 7.**
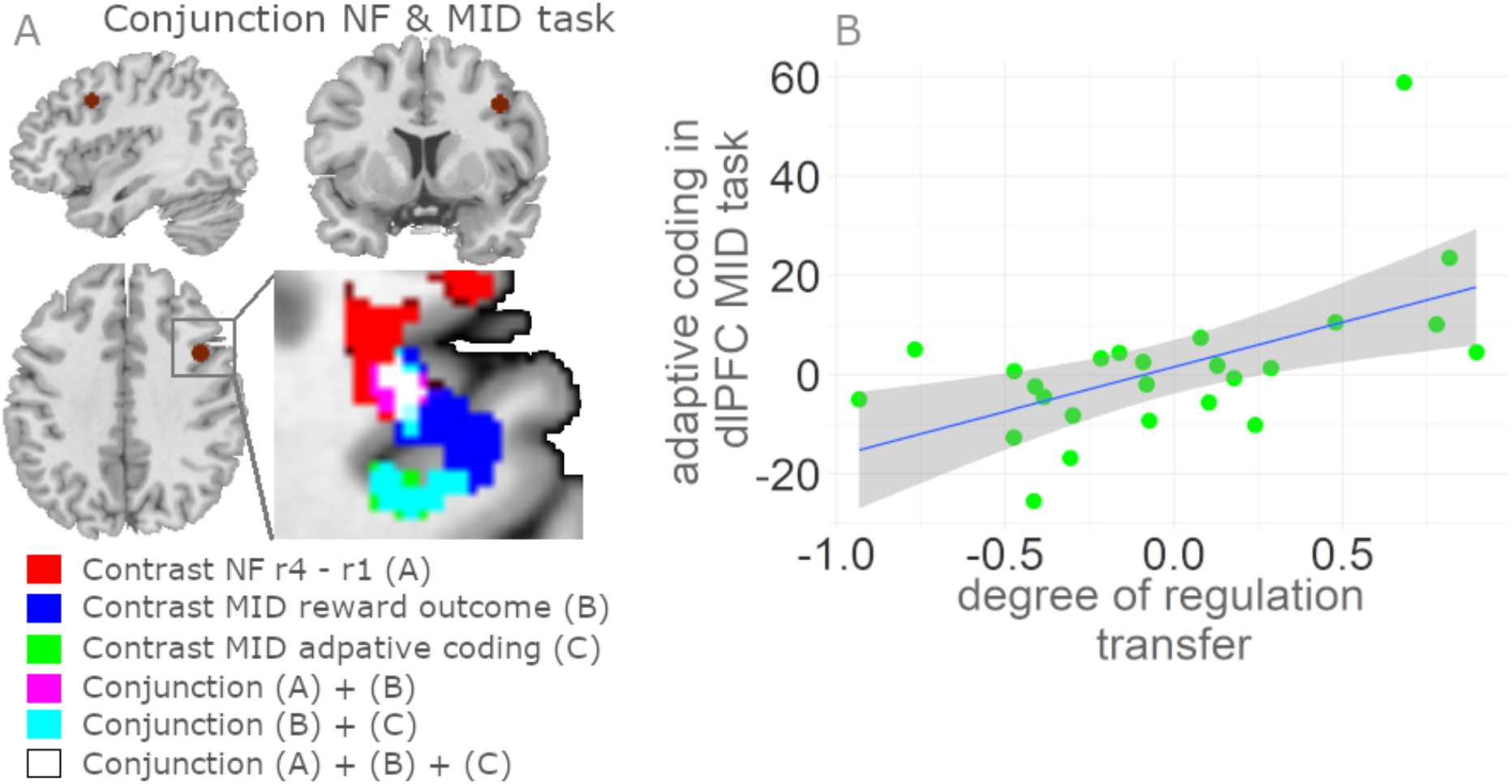
Reward-sensitivity in dlPFC correlates with successful SN/VTA self-regulation: (A) Degree of successful SN/VTA transfer (DRT) in the neurofeedback task correlated with prefrontal reward sensitivity and adaptive coding in the MID task. A conjunction analysis around the peak coordinate in dlPFC showing DRT-related decreases in prediction error coding during neurofeedback training (MNI x = 40, y = 10, z = 38, left) revealed common neural activity reflecting transfer (IMAGINE_REWARD_transfer_ − REST_transfer_) − (IMAGINE_REWARD_baseline_ − REST_baseline_) and reward sensitivity (small + large reward magnitude parametric modulators in MID, all contrasts with p<0.001). Moreover, individuals with more successful self-regulation of the SN/VTA showed stronger adaptive reward coding (which reflects higher sensitivity to small relative to large rewards) in the same region that also showed DRT-related decreases in prediction error coding during neurofeedback training (right). (B) The correlation plot depicts adaptive reward coding activity in dlPFC with DRT. The plot is for illustration purposes only without further significance testing to avoid double dipping. The grey shaded area identifies the 95 % confidence interval.

## 4 Discussion

In the present work, we used data acquired from two previous rt-fMRI neurofeedback studies to characterize individual differences and processes underlying successful transfer of self-regulation of the dopaminergic midbrain after neurofeedback training. This novel perspective on self-regulation success revealed insights on what distinguishes individuals who were more successful at SN/VTA regulation from those who were less successful. We found a significant relation between self-regulation success and increases in post-training activity in the cognitive control network. Moreover, we found four correlations with increasing transfer effects: (i) decreasing dlPFC prediction error signals during neurofeedback training, (ii) increasing connectivity of dlPFC with the SN/VTA for reward imagination compared to rest during transfer, (iii) increasing reward sensitivity in dlPFC and (iv) increasing adaptive reward coding in dlPFC in the independent MID task. Together, our study first shows that neurofeedback control of the dopaminergic midbrain relies on the cognitive control network. Second, our study suggests that the predictability of the upcoming feedback as reinforcement learning signal contributes to successful neurofeedback training.

Sustained self-regulation skills and the generalization of learning after neurofeedback training are key elements for practical applications and remain one of the major challenges in rt-fMRI neurofeedback research^49^. Results from previous neurofeedback studies of the reward system have been inconclusive^12,50,51^ and only one study^13^ reported significant post-training activity in the VTA, and increased mesolimbic network connectivity. Methodological limitations might have hampered the ability to detect transfer effects. First, previous studies focused exclusively on self-regulation of one *a priori* target region, such as SN/VTA, instead of investigating large-scale post-training effects within the whole brain. Second, transfer effects were examined at the group-level, which did not reflect the individual learning success. In the present study we overcome both limitations by taking advantage of an individual measure of transfer success (DRT) and focusing on the whole brain.

One insight of the present study is that transfer success associates with neural activity in cognitive control network areas^43,52^, such as dlPFC and ACC. The lack of cognitive control engagement within the control group and the correlation of DRT with the slope of SN/VTA increase during training in the intervention group only underpins that this finding is specific for the successful transfer of the learned self-regulation procedure. This network overlaps with regions that have been associated with feedback-related information processing during training^53,54^. Together, these findings suggest that the same regions contribute to acquisition and transfer of neurofeedback and that sustained post-training self-regulation generalizes across a functional network of different brain regions. Intriguingly, similar networks have been reported in skill learning. Future studies might investigate commonalities between neurofeedback and particularly cognitive skill learning, taking into account the specific temporal dynamics of both functions^22,55^.

The finding that individuals with more successful regulation of the dopaminergic midbrain show stronger activation of cognitive control areas during transfer speaks to our understanding of how individual differences in cognitive control affect emotion regulation^56–59^. For example the working memory component of cognitive control has been shown to predict negative affect reduction through reappraisal and suppression^60^. Interestingly, dopamine action (particularly at D1 receptors) in dlPFC sustains working memory performance^61^. Thus, it is conceivable that frontolimbic loops contribute to successful transfer. In any case, this notion converges with our finding of dlPFC-SN/VTA coupling being related to regulation success.

Future research might explore whether our findings, the positive post-training effects on the cognitive-control network activity, also have implications for transdiagnostic clinical applications. First, combining rt-fMRI neurofeedback training with different forms of psychotherapy such as cognitive behavioral therapy^62^, dialectical behavioral therapy^63^, or psychodynamic therapy^64–66^ could improve emotion regulation deficits prevalent in several psychiatric disorders including substance use disorders, depression, anxiety and personality disorders. It has already been shown that in patients suffering from depression, neurofeedback training can be a successful tool to re-stabilize modulation of the amygdala and increase its responsivity to reward^67^. It remains a question for future patient studies if a re-stabilization of cognitive control is also possible via such a training. With particular attention to substance use disorders, maladaptive changes in neuroplasticity within the cognitive control network are closely associated with loss of control and compulsive drug-seeking^68–70^. In these patients, neurofeedback training might be able to directly target the biological correlates and reinstate function of the cognitive-control network.

We found a reduction in prediction error coding in the DLPFC over the course of the neurofeedback training for successful regulators only, while these prediction error signals remained high for non-regulators. This finding suggests that prediction error-driven reinforcement learning was more pronounced in regulators than non-regulators and provides empirical evidence for previous theoretical proposals on the principles of neurofeedback learning independent of feedback modality^22^. Thus, reinforcement learning provides a framework for understanding how neurofeedback works. Future research may want to investigate whether the rich theoretical and empirical tradition of reinforcement learning^71^ can be harnessed to facilitate neurofeedback training.

We found that successful SN/VTA self-regulation is associated with an increased functional coupling between dlPFC regions coding prediction error and the dopaminergic midbrain. This coupling fits well with anatomical connections between dlPFC and the dopaminergic midbrain^19,21^ as well as effective connectivity studies on motivation^72^ and animal studies on prefrontal regulation of midbrain activity^20,73^. The animal work suggests that prefrontal cortex communicates with dopaminergic neurons primarily indirectly, through inhibitory relay neurons. By relating this coupling to successful midbrain self-regulation, our data go beyond previous connectivity studies of the dopamine system, which primarily focused on coupling between the prefrontal cortex and the striatum^74–76^.

At the functional level, a recent study on creative problem solving in humans highlights that dlPFC is involved in experiencing a moment of insight^77^. According to this effective connectivity study, dlPFC could upregulate the VTA/SN via striatal connections during such a moment. On the other hand, in trials where no solution was found for a given problem, also no significant connectivity was observed. This study supports our finding that dlPFC-SN/VTA connectivity plays an important role in self-guided motivation and in internal reward processing. Our finding points to the possibility that cognitive and affective mechanisms associated with different experiences also involve different neural pathways. Future studies should investigate to what degree individual differences in the functional architecture of brain networks^78^ influence these internal reward mechanisms and to which degree different strategies can influence neurofeedback training success.

Our independent reward task revealed that individual differences in prefrontal reward sensitivity and efficient adaptive reward coding were associated with successful SN/VTA self-regulation. Adaptive coding of rewards captures the notion that neural activity (output) should match the most likely inputs to maximize efficiency and representational precision^79^. Accordingly, we previously showed that reward regions encode a small range of rewards more sensitively than the large range of rewards^37,80^. Interestingly, in the present study, participants who were more sensitive to small rewards were also more successful in self-regulation of the dopaminergic midbrain. When participants in a typical neurofeedback training paradigm succeed at increasing the activity of the self-regulated area, the ensuing change in visual stimulation (positive neurofeedback) may constitute a small reward. By extension, adaptive reward coding may therefore provide a useful handle on identifying regulators. Moreover, future neurofeedback experiments should consider scaling the feedback signal to avoid sensitivity limitations, particularly in individuals with reduced adaptive coding.

A potential limitation of our study is that we used a combined mask for SN and VTA even though differences in functionality and anatomy have been reported for the two regions (reviewed e.g. by Trutti et al.^81^), with the SN more related to motor functions and the VTA to reward functions. However, it should be kept in mind that when viewed through the lens of recording and imaging rather than lesion techniques the differences are more gradual than categorical^82^. Still, future studies may want to use more specific feedback from one or the other region to more specifically target potential differences in functions. Further limitations are that only inverted feedback is available here as control group and this group has a smaller sample size. An additional control group perceiving no feedback could help to judge effects of neurofeedback training more precisely. Still, our data show a significant correlation between degree of regulation transfer and training runs only for the veridical feedback group and not for the control group. Moreover, it has been shown in other neurofeedback studies that volitional self-regulation of brain activity can only be learned when real feedback is presented^83^ and that other control groups failed to acquire VTA self-regulation^13^.

## 5 Conclusions

We showed that successful transfer in SN/VTA self-regulation after neurofeedback training is associated with activity in the cognitive control network (particularly dlPFC). Future studies could employ cognitive control activity during neurofeedback training to boost success rates and clinical outcomes. Furthermore, our findings of decreasing prediction error signals in dlPFC suggest that associative learning contributes to real-time fMRI neurofeedback effects. Finally, we show that higher individual reward sensitivity at the neural level increases the chance of neurofeedback training success. Patients with reduced neural reward sensitivity may therefore benefit from careful scaling of the neurofeedback information.

## Acknowledgments

The authors would like to thank Silvia Maier and Stephan Nebe for comments on previous versions of the manuscript and fruitful discussions. This project was supported by the European Union’s Horizon 2020 research and innovation program under the Grant Agreement No 794395 (to LH) and grant 100014_165884 from the Swiss National Science Foundation (to PNT). MK received grant support from the National Bank Fellowship (McGill) and Swiss National Science Foundation (P2SKP3_178175). The authors declare no competing financial interests.

