## Supplemental Material for "Individual differences in successful self-regulation of the dopaminergic midbrain"

### Supplemental Material on “Individual differences in mechanistic control of the dopaminergic midbrain”

### 1 Supplementary Methods

#### 1.1 MID task: Trial structure

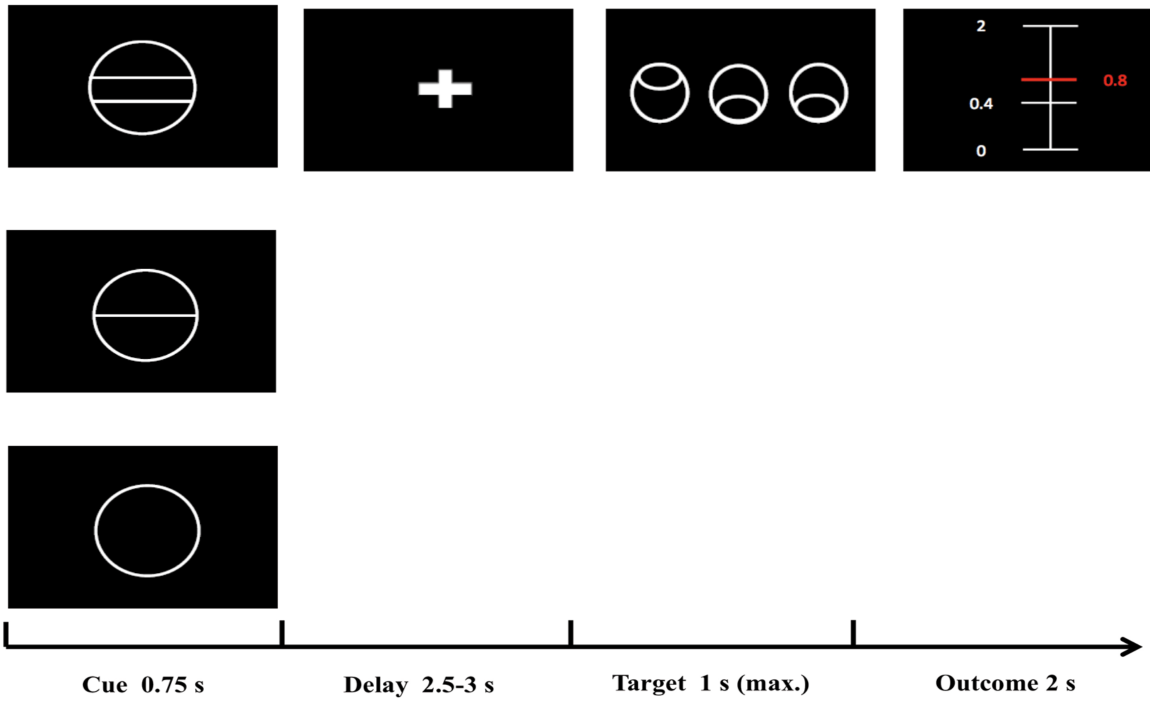

*Figure S 1* **Trial structure of the MID task:** First, one of three cues appeared. One cue was associated with large reward (ranging from 0 to 2.00 CHF), one cue with small reward (0 to 0.40 CHF) and one cue with no reward. After a delay of 2.5 to 3 s, participants had to identify an outlier from three circles by pressing one of three buttons as quickly as possible. Reward size depended on cue and response time.

### 2 Supplementary Results

#### 2.1 Cognitive control network analysis of correlation between degree of regulation transfer (DRT) and post-training effects in veridical feedback group

**Table S1**: Overlay of cognitive control template with regions showing significant correlation between DRT and [(IMAGINE_REWARD$_{transfer}$-REST$_{transfer}$) – (IMAGINE_REWARD$_{baseline}$ - REST$_{baseline}$)] in veridical group

| | | | MNI Coordinates | | |
|---|---|---|---|---|---|
| **Region Label** | **# voxels** | **t-value** | **x** | **y** | **z** |
| Middle Frontal Gyrus (dorsolateral prefrontal Cortex) | 436 | 4.61 | 45 | 31 | 19 |
| Left Thalamus | 536 | 4.47 | -9 | -24 | 10 |
| Temporal Occipital Fusiform Cortex | 701 | 5.85 | 32 | -46 | -22 |
| Occipital Fusiform Gyrus | 701 | 4.92 | 29 | -66 | -12 |
| Occipital Fusiform Gyrus | 562 | 4.87 | -42 | -70 | -19 |
| Right Cerebral White Matter (Right Middle Temporal Gyrus) | 122 | 4.95 | 56 | -45 | -4 |
| Left Caudate | 138 | 5.78 | -12 | 10 | 1 |
| Middle Frontal Gyrus | 151 | 5.35 | -39 | 5 | 53 |
| Right Cerebral White Matter (Insula) | 65 | 3.67 | 27 | 20 | -7 |
| Middle Frontal Gyrus | 90 | 4.54 | -42 | 17 | 40 |
| Right Thalamus | 160 | 4.40 | 5 | -13 | 13 |
| Left Cerebral White Matter | 152 | 4.30 | -45 | -45 | 1 |
| Frontal Pole | 145 | 4.15 | 24 | 59 | 7 |
| Precentral Gyrus | 169 | 4.07 | -41 | -9 | 62 |
| Lateral Occipital Cortex, superior division | 64 | 4.04 | 39 | -61 | 46 |
| Lateral Occipital Cortex, superior division | 109 | 3.96 | 27 | -64 | 52 |
| Lateral Occipital Cortex, superior division | 114 | 3.96 | -33 | -70 | 50 |
| Precuneus | 61 | 3.91 | 0 | -70 | 46 |
| Right Cerebral White Matter (dorsolateral prefrontal Cortex) | 89 | 3.88 | 36 | 16 | 37 |
| Frontal Orbital Cortex | 80 | 3.87 | -36 | 22 | -6 |
| Right Cerebral White Matter (Superior Temporal Gyrus) | 69 | 3.79 | 51 | -30 | 7 |
| Right Cerebral White Matter (Right Caudate Nucleus) | 68 | 3.76 | 14 | 10 | -1 |
| Right Cerebral White Matter (Superior Temporal Gyrus) | 69 | 3.79 | 51 | -30 | 7 |
| Right Cerebral White Matter (Right Caudate Nucleus) | 68 | 3.76 | 14 | 10 | -1 |
| Right Pallidum | 42 | 3.72 | 20 | -4 | 1 |

For all clusters within cognitive control template based on $t > 3.10$; $p < 0.001$; df = 40; minimum extent = 40; Table shows all local maxima separated by more than 20 mm. Regions were labeled using the Harvard-Oxford atlas and/or the Anatomy Toolbox in parentheses; the activity in SN/VTA has been excluded from the table to avoid circularity; x,y,z = Montreal Neurological Institute (MNI) coordinates in the left-right, anterior-posterior, and inferior-superior dimensions, respectively.

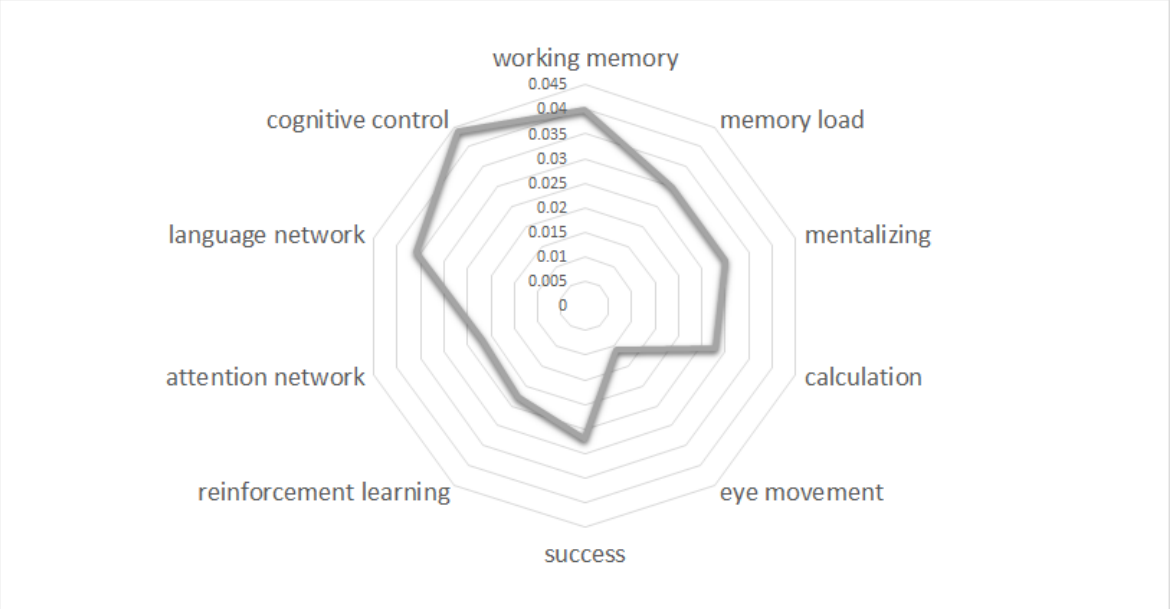

*Figure S 2* **Neurosynth database decoding**: To test the functional specificity of our results, we performed a meta-analytic functional decoding analysis using the Neurosynth database (neurosynth.org). We evaluated the representational similarity between the neural patterns of our cognitive control map. We selected nine different terms in domains covering task-related keywords. Results show that the neural signatures of the cognitive control decoding network reveal stronger similarity than other task-related neural patterns. Values on the spider plot represent Pearson's correlation coefficients.

#### 2.2 Correlation between DRT and post-training effects in inverted feedback group

**Table S2**: Regions showing significant correlation between DRT and [(IMAGINE_REWARD$_{transfer}$-REST$_{transfer}$) – (IMAGINE_REWARD$_{baseline}$ - REST$_{baseline}$)] in inverted feedback group

| | | | MNI Coordinates | | |
|---|---|---|---|---|---|
| **Region Label** | **# voxels** | **t-value** | **x** | **y** | **z** |
| Left Cerebral White Matter (Temporal Lobe) | 68 | 9.49 | -23 | -25 | -1 |
| Left Cerebral White Matter | 93 | 13.3 | -14 | -49 | 13 |
| Frontal Orbital Cortex (L IFG) | 71 | 8.38 | -29 | 16 | -21 |
| Left Thalamus | 332 | 7.69 | -3 | -24 | -4 |
| Right Thalamus | 332 | 6.10 | 18 | -27 | 1 |
| Left Cerebral White Matter (Frontal Lobe) | 367 | 7.65 | -29 | 13 | 19 |
| Left Cerebral White Matter (Frontal Lobe) | 174 | 7.09 | -18 | 46 | 5 |
| Left Amygdala | 51 | 6.34 | -26 | -4 | -21 |
| Left Cerebral White Matter (L IFG) | 125 | 6.33 | -42 | 29 | 8 |
| Location not in atlas (Temporal Lobe) | 173 | 6.03 | -5 | -13 | -10 |
| Insular Cortex | 56 | 6.02 | -45 | 5 | -9 |
| Lateral Occipital Cortex, superior division | 63 | 6.01 | -12 | -72 | 59 |
| Left Cerebral White Matter (Temporal Lobe) | 65 | 5.92 | -32 | 2 | -25 |
| Left Cerebral White Matter (Occipital Lobe) | 151 | 5.77 | -36 | -63 | 16 |
| Parahippocampal Gyrus, anterior division (L Fusiform Gyrus) | 137 | 5.72 | -36 | -12 | -27 |

| | | | | | |
|---|---|---|---|---|---|
| Left Cerebral White Matter (L Parahippocampal Gyrus) | 136 | 5.69 | -24 | -34 | -15 |
| Left Cerebral White Matter (Temporal Lobe) | 53 | 5.25 | -42 | -36 | 4 |
| Frontal Orbital Cortex (L IFG) | 63 | 5.03 | -33 | 26 | -12 |
| Superior Temporal Gyrus, anterior division | 114 | 4.98 | -54 | -4 | -10 |
| Left Cerebral White Matter (Temporal Lobe) | 56 | 4.83 | -45 | -39 | 8 |
| Temporal Pole | 50 | 4.44 | -39 | 1 | -43 |
| Inferior Temporal Gyrus | 43 | 4.58 | -48 | -40 | -19 |

For all clusters, $t > 3.70$; $p < 0.001$; df = 14; minimum extent = 40; Table shows all local maxima separated by more than 20 mm. Regions were labeled using the HarvardOxford atlas and/or the Anatomy Toolbox in parentheses; the activity in SN/VTA has been excluded from the table to avoid circularity; x,y,z =Montreal Neurological Institute (MNI) coordinates in the left-right, anterior-posterior, and inferior-superior dimensions, respectively.

#### 2.3 Disjunction analysis veridical and inverted feedback groups

**Table S3**: Disjunction analysis, reflecting preferential correlation between DRT and [(IMAGINE_REWARD$_{transfer}$-REST$_{transfer}$) – (IMAGINE_REWARD$_{baseline}$ - REST$_{baseline}$)] in veridical (Table S1) or control group (Table S2)

| | | | **MNI Coordinates** | | |
|---|---|---|---|---|---|
| **Region Label** | **# voxels** | **t-value** | **x** | **y** | **z** |
| Left Cerebral White Matter (L Precuneus) | 87 | 13.31** | -14 | -49 | 13 |
| Left Cerebral White Matter (Temporal Lobe) | 62 | 9.49** | -23 | -25 | -1 |
| Frontal Orbital Cortex (L IFG) | 64 | 8.38** | -29 | 16 | -21 |
| Left Thalamus | 257 | 7.69** | -3 | -24 | -4 |
| Right Thalamus | 257 | 6.10** | 18 | -27 | 1 |
| Left Cerebral White Matter (Frontal Lobe) | 333 | 7.65 | -29 | 13 | 19 |
| Left Cerebral White Matter (Frontal Lobe) | 88 | 7.09 | -18 | 46 | 5 |
| Left Amygdala | 48 | 6.34** | -26 | -4 | -21 |
| Left Cerebral White Matter (L IFG) | 91 | 6.33** | -42 | 29 | 8 |
| Insular Cortex (Temporal Pole) | 48 | 6.02** | -45 | 5 | -9 |
| Lateral Occipital Cortex, superior division | 55 | 6.01 | -12 | -72 | 59 |
| Left Cerebral White Matter (Temporal Lobe) | 57 | 5.3 | -32 | 2 | -25 |
| Left Cerebral White Matter (Occipital Lobe) | 132 | 5.77 | -36 | -63 | 16 |
| Parahippocampal Gyrus, anterior division | 122 | 5.72** | -36 | -12 | -27 |
| Left Cerebral White Matter (Frontal Lobe) | 65 | 5.54 | -9 | 37 | 8 |
| Frontal Orbital Cortex (L IFG) | 54 | 5.03** | -33 | 26 | -12 |

| | | | | | |
|---|---|---|---|---|---|
| Superior Temporal Gyrus | 87 | 4.98** | -54 | -4 | -10 |
| Temporal Pole | 42 | 4.43 | -39 | 1 | -43 |

For all clusters, t > 3.10; p < 0.001; minimum extent = 40; Table shows all local maxima separated by more than 20 mm. Regions were labeled using the HarvardOxford atlas and/or the Anatomy Toolbox in parentheses; x,y,z =Montreal Neurological Institute (MNI) coordinates in the left-right, anterior-posterior, and inferior-superior dimensions, respectively.
** Significant difference in direct comparison between veridical and inverted feedback groups (p <.001).

#### 2.4 Conjunction analysis of veridical and inverted feedback groups

**Table S4**: Conjunction analysis, reflecting common correlation between DRT and [(IMAGINE_REWARD$_{transfer}$-REST$_{transfer}$) – (IMAGINE_REWARD$_{baseline}$ - REST$_{baseline}$)] in veridical (Table S1) and inverted feedback group (Table S2)

| | | | MNI Coordinates | | |
|---|---|---|---|---|---|
| **Region Label** | **# voxels** | **t-value** | **x** | **y** | **z** |
| Temporal Occipital Fusiform Cortex | 28 | 5.81 | 32 | -46 | -24 |
| Right Cerebral White Matter | 154 | 5.80 | 5 | -16 | -10 |
| Right Cerebral White Matter (Right Hippocampus) | 32 | 5.26 | 21 | -18 | -10 |
| Left Thalamus | 45 | 5.1 | -8 | -16 | -1 |
| Inferior Temporal Gyrus, posterior division | 85 | 4.20 | -60 | -40 | -16 |
| Parahippocampal Gyrus (Fusiform Gyrus) | 32 | 3.93 | -21 | -34 | -18 |
| Parahippocampal Gyrus, posterior division | 25 | 3.86 | -15 | -45 | -16 |

For all clusters, t > 3.10; p < 0.001; minimum extent = 20 (due to conjunction of two contrasts); Table shows all local maxima separated by more than 20 mm. Regions were labeled using the HarvardOxford atlas and/or the Anatomy Toolbox in parentheses; x,y,z =Montreal Neurological Institute (MNI) coordinates in the left-right, anterior-posterior, and inferior-superior dimensions, respectively.

#### 2.5 Decreasing prediction error coding

**Table S5**: Cluster table for prediction error coding analysis during neurofeedback training based on contrast (IMAGINE_REWARD-REST)training_run2 – (IMAGINE_REWARD-REST)training_run1

| | | | MNI Coordinates | | |
|---|---|---|---|---|---|
| **Region Label** | **# voxels** | **t-value** | **x** | **y** | **z** |
| Lateral Occipital Cortex, inferior division | 197 | 4.847 | -45 | -69 | -10 |
| Postcentral Gyrus | 210 | 4.511 | 50 | -25 | 44 |
| Right Cerebral White Matter (dorsolateral prefrontal cortex) | 23 | 3.393 | 35 | 11 | 37 |
| Right Cerebral White Matter | 49 | 4.357 | 30 | -21 | 49 |
| Superior Temporal Gyrus, posterior division | 36 | 4.241 | -68 | -40 | 13 |
| Left Cerebral White Matter (Left Inferior Frontal Gyrus) | 55 | 4.083 | -42 | -57 | -3 |
| Left Cerebral White Matter (Left Caudate Nucleus) | 31 | 4.072 | -18 | -4 | 28 |

| | | | | | |
|---|---|---|---|---|---|
| Right Cerebral White Matter (Right Inferior Occipital gyrus) | 39 | 4.068 | 39 | -73 | -3 |
| Postcentral Gyrus | 68 | 4.027 | -41 | -27 | 49 |
| Superior Parietal Lobule | 29 | 4.010 | 14 | -51 | 71 |
| Right Cerebral White Matter | 30 | 3.972 | 39 | -37 | 31 |
| Superior Parietal Lobule | 53 | 3.862 | -39 | -40 | 47 |
| Right Cerebral White Matter | 36 | 3.838 | 29 | -48 | 46 |
| Precentral Gyrus | 26 | 3.827 | 0 | -22 | 53 |
| Left Cerebral White Matter | 34 | 3.820 | -12 | -88 | 8 |
| Supracalcarine Cortex | 61 | 3.758 | 24 | -61 | 19 |
| Juxtapositional Lobule Cortex (formerly Supplementary Motor Cortex) | 21 | 3.758 | -3 | -3 | 47 |
| Lateral Occipital Cortex (Precuneus) | 56 | 3.751 | 14 | -61 | 59 |
| Lateral Occipital Cortex | 72 | 3.561 | -20 | -73 | 52 |
| Lateral Occipital Cortex | 24 | 3.559 | -26 | -76 | 38 |

For all clusters, t > 3.1; p < 0.001; df = 40; minimum extent = 20; Table shows all local maxima separated by more than 20 mm. Regions were labeled using the HarvardOxford atlas and/or the Anatomy Toolbox in parentheses. The activity in SN/VTA has been excluded from the table; x,y,z =Montreal Neurological Institute (MNI) coordinates in the left-right, anterior-posterior, and inferior-superior dimensions, respectively.

#### 2.6 Reward adaptation and reward sensitivity in MID task correlated with DRT

**Table S6**: Cluster table for correlation of DRT with MID task-based reward adaptation (difference of small minus large reward parametric modulators) and reward sensitivity (sum of small and large reward parametric modulators) as conjunction of all three analysis

| | | | **MNI Coordinates** | | |
|---|---|---|---|---|---|
| **Region Label** | **# voxels** | **t-value** | **x** | **y** | **z** |
| Middle Frontal Gyrus (dorsolateral prefrontal cortex) | 21 | 3.72 | 39 | 8 | 39 |
| Temporal Occipital Fusiform Cortex | 37 | 4.98 | 32 | -43 | -21 |
| Middle Temporal Gyrus, temporooccipital part | 344 | 4.60 | -63 | -48 | -4 |
| Left Cerebral White Matter | 32 | 4.30 | -45 | -45 | 1 |
| Frontal Orbital Cortex | 64 | 4.27 | -51 | 26 | -13 |
| Right Thalamus | 24 | 4.77 | 18 | -28 | -4 |
| Left Cerebral White Matter | 23 | 4.12 | -8 | -16 | -6 |
| Superior Frontal Gyrus | 38 | 3.86 | -2 | 37 | 58 |
| Parahippocampal Gyrus | 22 | 3.71 | 20 | -12 | -28 |

For all clusters, t > 3.1; p < 0.001; df = 40; minimum extent = 20 (due to conjunction); Table shows all local maxima separated by more than 20 mm. Regions were labeled using the HarvardOxford atlas and /or the AnatomyToolbox in parentheses; x,y,z =Montreal Neurological Institute (MNI) coordinates in the left-right, anterior-posterior, and inferior-superior dimensions, respectively.

#### 2.7 Strategies used

**Table S7:** Overview of the proposed strategies and how often these strategies had been chosen by the participants

| Proposed strategy used by | Veridical feedback Group (N=42) | Inverted feedback Group (N = 17) |
|---|---|---|
| **Family and friends** | 27 | 3 |
| **Food** | 7 | 1 |
| **Personal achievement** | 9 | 1 |
| **Romantic or sexual imagery** | 39 | 8 |
| **Leisure activities** | 9 | 5 |
| **Individual others** | - | - |

#### 2.8 Prediction error coding in early and late neurofeedback training phase

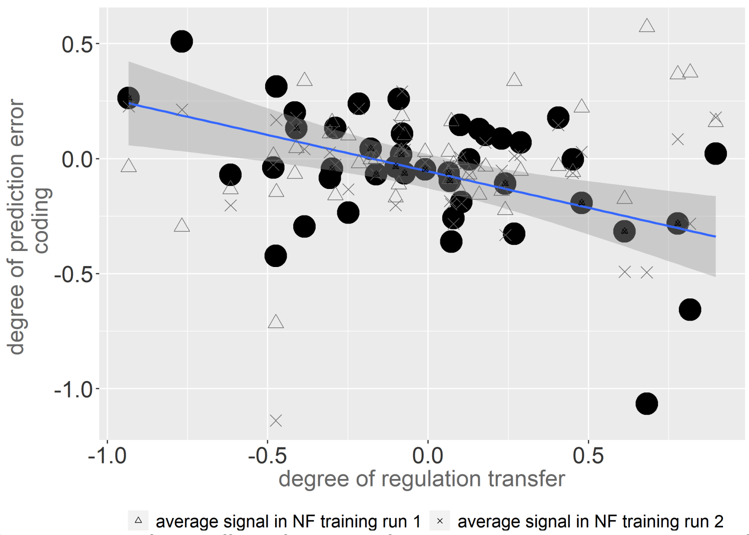

*Figure S3*: **Prediction error coding in dlPFC decreases during NF training in participants with successful SN/VTA self-regulation:** The time-resolved prediction error signal in dlPFC, corresponding to the parametric difference between the current and immediately preceding feedback activity from the SN/VTA decreased with ongoing feedback training for the successful participants only. Here, the difference between run 2 and run 1 is replotted from Figure 5. Moreover, the contrast estimate from the parametric prediction error modulator is illustrated separately for runs 1 and 2, showing decrease of prediction error in late training for successful regulators.

#### 2.9 Control analysis for spatial specificity of dopaminergic midbrain regulation

To investigate the spatial specificity of our analysis of dopaminergic midbrain regulation, we performed the same analysis as described in the main text for SN/VTA but using the neighboring brain region of the parahippocampus as control ROI. This target area is also active during the self-regulation task because the participants perform memory-based strategies. We extracted the parahippocampus mask from the human Talairach atlas in the WFU Pickatlas and performed the identical main effects analysis as described in section 2.5 of the manuscript. To assess commonalities for the two target ROIs, SN/VTA and parahippocampus, we performed a conjunction analysis based on the unthresholded contrast image of the parahippocampus. This analysis revealed two common areas within the cerebellum and the temporal gyrus (Figure S4 and Table S8). The limited commonalities between these two target ROIs, especially in striatal and prefrontal areas, indicate spatial specificity of our findings using the SN/VTA as target region. To corroborate this interpretation, we also directly compared the results of the two target ROIs in a second level analysis (SN/VTA - parahippocampus) and calculated the conjunction of this difference with the SN/VTA results only. The results of this conjunction analysis revealed similar neural activity patterns as the SN/VTA pattern only. This analysis underpins the local specificity of our results because the interpretation of the results remains the same even when the parahippocampus result pattern has been subtracted.

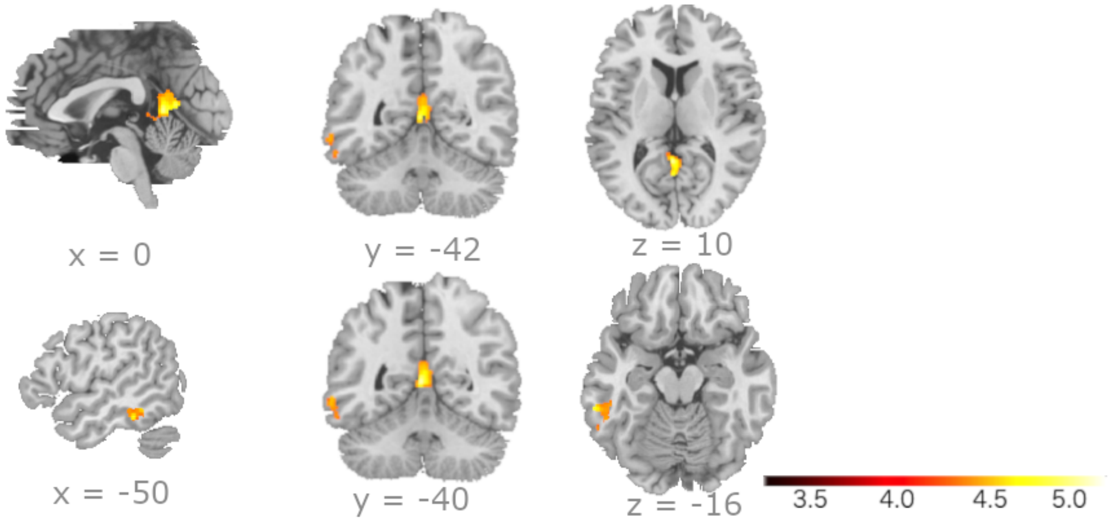

*Figure S4:* **Spatial specificity of SN/VTA findings:** Conjunction analysis of two different ROIs – our target ROI SN/VTA and control ROI in parahippocampus – for the main effect analysis of individual regulation transfer revealed limited commonalities between these two target ROIs, especially in striatal and prefrontal areas. This is in keeping with spatial specificity of our SN/VTA findings.

**Table S8:** Clusters overlapping in main (coactivation with SN/VTA) and control (coactivation with parahippocampus) analysis

| | | | MNI Coordinates | | |
|---|---|---|---|---|---|
| **Region Label** | **# voxels** | **t-value** | **x** | **y** | **z** |
| Cerebellar Vermis (4/5) | 701 | 6.104 | -3 | -49 | 7 |
| L Inferior Temporal Gyrus | 326 | 4.913 | -62 | -37 | -16 |

For all clusters, $t > 3.10$; $p < 0.001$; df = 40; minimum extent = 20 (due to conjunction); Table shows all local maxima separated by more than 20 mm. Regions were labeled using the HarvardOxford atlas and /or the AnatomyToolbox in parentheses; x,y,z =Montreal Neurological Institute (MNI) coordinates in the left-right, anterior-posterior, and inferior-superior dimensions, respectively.

#### 2.10 Distribution of DRT

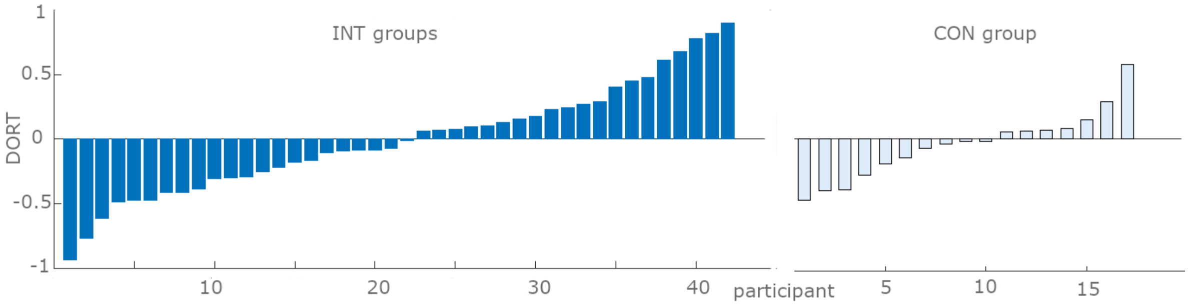

*Figure S5: Distribution of DRT measure in intervention (INT) groups and control (CON) group*
